# Increasing the efficacy of exposure using a whole brain approach to real-time fMRI neurofeedback among female victims of interpersonal violence

**DOI:** 10.1101/2022.08.19.504571

**Authors:** Maegan L. Calvert, Josh M. Cisler, Keith A. Bush

## Abstract

Individuals who experience interpersonal violence (IPV) and poly-victimization have an increased likelihood of developing Posttraumatic Stress Disorder (PTSD), and statistically, women are more likely than men to be victims of IPV, experience poly-victimization, and develop PTSD. Current gold-standard PTSD treatments utilize exposure, a key mechanism of change; however, exposure-based treatments suffer from moderate remission rates. This outcome underscores the importance of utilizing knowledge of neural mechanisms to increase treatment effectiveness and PTSD remission rates. The current study used a symptom provocation paradigm, which is conceptually similar to exposure, and whole brain multivariate pattern classifiers (MVPC) to provide participants with neurofeedback on their ability to regulate their brain-states. In the MRI scanner, women (*N* = 21; control *n* = 11, PTSD *n* = 10) completed a symptom provocation paradigm. The MVPC was constructed during the first two runs and neurofeedback was given during the third and fourth run. During all four runs, participants were instructed to increase or decrease their emotional engagement with their stress/trauma script and during the last two runs, participants were provided visual feedback indicating their performance in regulating brain states. Skin conductance response was utilized as an independent measure of physiological arousal. Generalized linear models indicated a significant interaction effect of feedback and condition as well as feedback and group. Generalized linear models of skin conductance response largely mirrored these findings. Results indicate neurofeedback of whole brain activation can be utilized to increase engagement with trauma memories. Thus, exposure-based treatments, possibly with refractory cases of PTSD, may be improved with neurofeedback by providing the therapist and patient insight into their brain-state during exposure which may increase the effectiveness of exposure-based treatments.

---

Interpersonal violence (i.e., child maltreatment, sexual assault, intimate partner violence) affects women at a disproportionate rate as compared to men^1^, and women who experience interpersonal violence have an increased likelihood of developing post-traumatic stress disorder (PTSD ^2–4^). PTSD is characterized by heterogeneous symptoms including hyperarousal, intrusive memories of the experience, avoidance of reminders, and negative alterations in cognitions^5^ which are subsequently associated with interpersonal conflict, emotion dysregulation, and alexithymia^6^. Among women who experience interpersonal violence, the prevalence of PTSD ranges from 31% ^6^ to 70% ^7^. Because PTSD causes significant difficulties in living, there has been a proliferation in empirically supported treatments designed to address PTSD symptoms^8–10^.

One core component of PTSD treatments is exposure and it is often considered a primary mechanism in the reduction of symptoms^9^. Exposure involves an individual repeatedly experiencing a feared stimulus in a safe environment as a means for the individual to learn that the feared stimulus is no longer a signal for danger^11^. Specifically, an individual suffering from PTSD often experiences significant distress when they encounter a stimulus that reminds them of the trauma memory^5^. For example, a woman who was sexually assaulted and now experiences symptoms consistent with PTSD might encounter a man who resembles the attacker. This encounter might trigger the memory of the sexual assault resulting in a cascade of emotional and behavioral dysregulation. An individual with PTSD may also avoid thoughts, feelings, and reminders of the trauma which may result in avoidance of close relationships, intimacy, or people, places, and things that are important to the individual’s daily life^5^. An exposure-based intervention involves repeated confrontation of the trauma memory, through retelling the event narrative, to weaken the avoidance related to or emotional distress elicited by the trauma memory and enable exposure to corrective information^12,13^. While exposure-based treatments are the psychological treatment of choice for PTSD and meta-analyses suggest large effect sizes ^14^, these treatments are associated with post-treatment remission rates of only ~53-65% ^8,15^. Moreover, among individuals whose PTSD symptoms remit after treatment, ~20% report non-clinically meaningful change in self-reported social functioning and ~50% report non-clinically meaningful change in self-reported quality of life ^15^. Given the moderate PTSD remission rates and quality of life improvement, basic and clinical scientists need a more thorough understanding of the brain’s response to treatment and the underlying neural mechanisms that mediate PTSD symptomology in order to support the delivery of more effective treatment.

There are different ways in which scientists have sought to understand the neural mechanisms of PTSD and responses to treatment: a) correlational studies after treatment, b) experimental paradigms that induce symptoms and record brain responses to trauma reminders, and c) experimental paradigms that induce symptoms and record brain responses while also instructing participants to increase or decrease a given brain-state by providing real-time brain-state feedback. We will briefly review correlational and experimental paradigms without feedback to provide context, and then more fully review the experimental real-time feedback studies in the context of PTSD. Because exposure is the most common mechanism of symptom change in current PTSD treatments, we will only review findings related to symptom provocation paradigms which are conceptually similar to exposure in behavioral and cognitive-behavioral treatment approaches.

Existing correlational and experimental studies support the hypothesis that psychological treatments alter neural mechanisms and mediate core PTSD symptoms. For example, in a review of pre- and post-treatment neural correlate studies, reductions in PTSD symptoms were consistently associated with reductions in hemodynamic activity from pre-treatment to post-treatment in both the amygdala and insula ^16^. Symptom improvement was also associated with increases in hemodynamic activity from pre-treatment to post-treatment in the dorsal and ventral anterior cingulate cortex, hippocampus, and prefrontal cortex ^16^.

In experimental symptom provocation studies, participants recount details of their trauma experiences to a trained research specialist prior to the fMRI scan. Research specialists then transform this information into stimuli that can be utilized in the MRI environment. Participants view or hear trauma reminders (e.g., narrative script or key words) while in the MRI scanner which is then followed by neutral stimuli or a rest period ^17^. Experimental paradigms using trauma-related stimuli have varying results depending on paradigm and analysis type. In a recent review of symptom provocation studies, the authors concluded studies consistently reported hyperactivation in the dorsal anterior cingulate cortex (DACC) and hypoactivation in the ventromedial prefrontal cortex (vmPFC)^17^. These authors also indicated that there were common brain regions recruited during symptom provocation studies such as the amygdala, hippocampus, rostral anterior cingulate cortex, and insula, but that the direction of these associations varied by analysis-type and comparison group-type ^17^.

In order to more fully understand the neural mechanisms of PTSD symptom induction and mechanisms of possible symptom reduction/remission, scientists have utilized real-time functional magnetic resonance imaging (rt-fMRI) and neurofeedback. Real-time fMRI refers to near-instantaneous recording and analyzing of fMRI signal while participants are inside the scanner. This provides an opportunity to present information (typically visually) regarding a participant’s brain-state immediately to the participant. Using this information, the participants are instructed to modulate (i.e., self-regulate) their brain-state in desirable ways ^18–20^. Real-time fMRI neurofeedback (rt-fMRI-nf) paradigms have been utilized in healthy controls ^21,22^ and in patients with symptoms of depression ^23,24^, anxiety ^25,26^, and PTSD ^27–35^.

However, fewer studies have utilized symptom provocation experiments during rt-fMRI-nf paradigms ^27,28,31,32,35^. These studies utilize symptom provocation to generate PTSD-related brain-states and then participants are tasked with regulating their brain-states with the help of feedback. Previous work suggests that brain-state modulation can induce changes in the amygdala and other brain regions after neurofeedback training. For example, when using amygdala down regulation rt-fMRI-nf, participants demonstrated significantly more activation in the dorsal lateral prefrontal cortex (dlPFC) and right ventrolateral prefrontal cortex (vlPFC) ^31^ during blocks where participants regulated their amygdala activity as compared with blocks where participants passively viewed the provocation stimuli. In addition, networks hypothesized to play a significant role in PTSD (i.e., central executive network, salience network, and default mode network) appear to be differentially modulated by amygdala down-regulation rt-fMRI-nf training sessions ^32^. In addition to immediate brain changes, participants also evidence increased connectivity between the amygdala and regulatory regions one week after amygdala down-regulation rt-fMRI-nf ^28^.

Although studies largely focus on amygdala down-regulation for PTSD, there is a small, but growing literature that this approach may be limited. In general, studies suggest that rt-neurofeedback may engage many different brain areas rather than specific regions of interest (ROIs)^29,36,37^. For rt-fMRI-nf without regard to diagnosis/problem, studies demonstrate that neurofeedback training activates brain regions related to self-awareness like the insula and higher level functions like the anterior cingulate cortex (ACC) and ventrolateral prefrontal cortex (vlPFC) ^36,37^. In regard to rt-fMRI-nf for PTSD specifically, multivariate pattern classification and predictions are most reliable when using whole brain activations rather than ROIs^27^. Similarly, findings also indicate that brain regions involved in self-awareness and memory moderate associations among PTSD brain-states and symptom decreases ^29^. Consequently, the current state of the literature regarding the neural mechanisms of PTSD and symptom reduction/maintenance is not able to address whole brain interactions or speak to utilizing the whole brain in order to increase treatment effectiveness.

## Objective

Given this context, we aimed to 1) utilize a whole brain rt-fMRI-nf symptom provocation paradigm to determine if whole brain activations could be used to modulate trauma-related brain activity and 2) sought to determine if this approach would be commensurate with objective measures of arousal and disengagement.

## Hypothesis

1. Real-time fMRI neurofeedback of whole brain activations can be used to modulate trauma-related brain states.
2. Increases or decreases of whole brain activity will correspond to instructions to engage or disengage with the trauma script.
3. Increases and decreases in whole brain activity will correspond to an objective convergent measure of arousal (i.e., skin conductance response).

## Method

### Ethics Statement

All participants provided written informed consent after receiving written and verbal descriptions of the study procedures, risks, and benefits. We performed all study procedures and analysis with approval and oversight of the Institutional Review Board at the University of Arkansas for Medical Sciences (UAMS) in accordance with the Declaration of Helsinki and relevant institutional guidelines and policies.

### Participants and Assessment

Women (*N* = 21) age 21-50 years (*M* = 32.95, *SD* = 9.33) were recruited into the current study from outpatient mental health clinics and community wide advertisements. Participants were classified as 1) minimally symptomatic controls with or without a history of assault or 2) PTSD positive (i.e., meeting full criteria or just sub-threshold for PTSD diagnosis) with a history of physical or sexual assault. Exclusion criteria consisted of severe current medical condition, psychotic disorders, or internal ferromagnetic objects. All study procedures were approved by the University of Arkansas for Medical Sciences institutional review board and all participants provided written informed consent. This experiment was preregistered on Clinicaltrials.gov [NCT02500719], and results were uploaded on August 20, 2021.

All participants attended two assessment sessions. During the first session, participants completed the Structured Clinical Interview for DSM-IV Disorders^38^ administered by a trained clinical interviewer to assess for lifetime and current DSM diagnoses, as well as, the PTSD Checklist for DSM-5 (PCL-5^39^) to assess for severity of PTSD symptoms. Participants also completed a stress/trauma script (see below). During the second session, participants engaged in fMRI procedures.

In the control group (n = 11), participants (*M* = 30.09 years, *SD* = 8.49) identified as White (82%, n = 9) and Black (18%, n=2). In the PTSD group (n = 10), participants (*M* = 36.10 years, *SD* =9.61) identified as White (70%, n = 7), Asian (n = 1), Black (n = 1), and Latino (n = 1). On the SCID-IV, 55% of the control group (n = 6) and 100% of the PTSD group (n = 10) had a lifetime or current diagnosis other than PTSD. Women in the control group had significantly less PTSD symptoms (*M* = 10.64, *SD* = 11.16) than women in the PTSD group (*M* = 48.60, *SD* = 11.24,*p* < .001). See supplemental table S1.

### Script Tasks

Participants in the control condition generated a stress script to activate a stress memory while participants in the PTSD condition generated a trauma memory script to activate the trauma memory. Consistent with previous literature, participants were instructed to describe the event, contextual information that occurred before and after the event as well as bodily sensations that occurred during the event^40–43^. These short narratives were approximately 150-200 words and were displayed on a screen while participants were in the MRI scanner. Verbal narrations of the scripts were also recorded in the voice of a female research assistant. The scripts were read aloud such that the total narration lasted approximately one minute and were played through the participants’ headphones during the scan.

### MRI Acquisition

All imaging data were acquired via a Philips 3T Achieva X-series MRI scanner (Philips Healthcare, Eindhoven, The Netherlands) with a 32-channel head coil. Anatomic images were acquired using an MPRAGE sequence (matrix = 256 x 256, 220 sagittal slices, TR/TE/FA = 8.0844/3.7010/8°, final resolution =0.94 x 0.94 x 1 mm^3^). Functional images were acquired using the following EPI sequence parameters: TR/TE/FA = 2000 ms/30 ms/90°, FOV = 240 x 240 mm, matrix = 80 x 80, 37 oblique slices, ascending sequential slice acquisition, slice thickness = 2.5 mm with 0.5 mm gap, final resolution 3.0 x 3.0 x 3.0 mm^3^.

MR imaging commenced with a T1-weighted structural scan followed by four functional MRI (fMRI) acquisition runs. The first two fMRI acquisition runs consisted of repeated 150 s block presentations of the stress/trauma script. These data were used to train the machine learning model to be used in real-time fMRI acquisition (see below). Script presentations were preceded by instructions either to engage with the trauma/stress script to increase arousal or to disengage with the script to suppress arousal. The order of engagement and disengagement was counterbalanced across the two runs. Prior to scanning, participants selected the engagement and disengagement strategies they would use from a menu of possible strategies (e.g., focusing on the worst part of the trauma/stress memory to engage or focusing on breathing to disengage).

The final two runs of the fMRI acquisition consisted of repeated 20 s block presentations of the stress/trauma script preceded by instructions to either engage or disengage with the script to increase or decrease arousal. Half of these blocks provided neurofeedback (by varying the transparency of the script’s text relative to the background) to inform participants how successfully they were engaging or disengaging with the script based upon real-time predictions of the machine learning model. The remaining blocks presented the script’s text at a fixed level of 50% transparency throughout.

### Skin Conductance Data Acquisition

Skin conductance was concurrently recorded alongside MRI using the BIOPAC (Goleta, CA) MP150 Data Acquisition System and AcqKnowledge software combined with the EDA100C-MRI skin conductance acquisition module. Following prior reported work^44^, recording electrodes were placed on the medial portions of the thenar and hypothenar eminences of the left hand; a ground electrode was placed on the ventral surface of the left wrist.

### Data Analysis

#### Real-time MR Image Processing, Multivariate Pattern Classification, and Neurofeedback

We implemented custom computer code that acquired each raw fMRI volume as it was written to disk (post-reconstruction) and, using AFNI^45^, performed motion correction via rigid body alignment (corrected to the tenth volume of the first run), incremental detrending (with mean preserved) based upon the set of volumes acquired to that point in the scan, spatial smoothing using a 8 mm FWHM Gaussian filter, and segmentation into gray matter (GM), white matter, and cerebrospinal fluid.

A support vector machine multivariate classifier, implemented via the LIBSVM library^46^ was fit to whole-brain GM-masked fMRI data drawn from the first two runs. Volumes acquired during script engagement and disengagement task blocks were assigned class labels +1 and −1, respectively. Volumes exceeding motion limits (FD>0.5) were excluded from training. The SVM used a radial basis function kernel and a cost parameter of 100 derived from earlier work on similar task data^27^.

The trained classifier was then applied to predict the class label of each volume acquired during the final two fMRI acquisition runs. These predictions took the form of Euclidean distances from a decision hyperplane which separates fMRI volumes containing patterns of neural activation that resemble engaged training examples (positive distances) in comparison to disengaged examples (negative distances). During neurofeedback blocks, predictions of hyperplane distance were converted into levels of transparency of the trauma/stress script’s text to provide participants with real-time visual feedback of their task performance. Volumes with increasingly positive hyperplane predictions (i.e., classified as increasingly emotionally engaged) generated increasingly opaque yellow text of the trauma/stress script against the static black background. That is, as the classifier predicted they were more emotionally engaged with the trauma/stress memory, the text of the script grew brighter. Volumes with increasingly negative hyperplane predictions (i.e., classified as decreasingly emotionally engaged) generated increasingly transparent text of the trauma/stress script against the static black background. That is, as the classifier predicted they were less emotionally engaged with the trauma/stress memory, the text of the script faded from view. Transparency values were interpolated between classifier predictions at 20 Hz to produce smoothly varying text transparency despite the acquisition of fMRI volumes every 2 s. Both generalized linear models and linear mixed effects models (which account for nested observations within individuals and individual variability) were utilized to assess the effect of experimental manipulations on hyperplane distances. In all cases, the mixed effects linear models provided the better fit according to AIC, BIC, and Log Likelihoods^47^ and will be presented in the results. To interpret the interactions in the linear mixed effects models, we calculated the difference between the estimates and the pooled standard error, and then calculated the t-value, degrees of freedom, and the respective p-value for each comparison.^48^ Exploratory analyses for post-hoc voxel-wise comparisons using linear mixed effects were conducted to visualize the contributions of the attention networks in the feedback conditions while engaging and disengaging in the trauma/stress script and are described in the Supplemental Materials.

#### Skin Conductance Response Processing

Skin conductance was acquired at 2000 Hz. These data were transformed according to best practices^49–51^ as follows: 20 ms median filtered, subsampled to 200Hz, zero-meaned, and then both low- (5 Hz) and high-pass (0.0159 Hz) filtered. The resulting data were then z-scored within each run. We then implemented a general linear convolution model-based regression (GLCM ^49,50,52–54^) to estimate the skin conductance response (SCR) within each volume of fMRI acquisition as it has evidenced sensitivity in previous studies.

## Results

### Whole Brain Activity in Response to Neurofeedback

A mixed effects linear model (see Tables 1 and 2) was used to assess the effect of condition (CDN; instructions to engage or disengage from the trauma/stress script), neurofeedback (FB; feedback given or not given), and group membership (GRP; control or PTSD) on whole brain activity as measured by hyperplane distances. Main effects of CDN, FB, RUN, and GRP were not significant. However, interaction effects for CDN x FB (fixed effect: r = 0.22, *p* = 0.000067, Wald *t*, null: β = 0) and CDN x GRP (fixed effect: r = 0.38, *p* = 0.04, Wald *t*, null: β = 0) were significant when accounting for random effects. Simple effects models indicated hyperplane distances were significant for the engage condition with feedback (fixed effect: r = 0.07, *p* = 0.00, z = 1.91, null: β = 0) but not the disengage condition with feedback (fixed effect: r = −0.02, *p* = 0.60, *z* = −0.48, null: β = 0); however, the difference between the two effects was significant [*t* (7,093) = 6.11, *p* = .00]. Hyperplane distances were significant for both groups in the engage condition, but the PTSD group had larger hyperplane distances (fixed effect: r = 0.31, *p* = 0.00, z = 15.56, null: β = 0) than the control group (fixed effect: r = 0.10, *p* = 0.00, z = 7.96, null: β = 0) which was significant [*t* (7,093) = −2.91, *p* = .00]. Models also demonstrated that hyperplane distances were not significant for either group in the disengage condition (fixed effect: r = 0.14, *p* = 0.14, z = 1.50, null: β = 0). This mixed effects linear model accounted for 27% of the variance in hyperplane distances (*R^2^* = 0.27). For all simple effect comparisons see Table 2, for visualizations of changes in hyperplane distances TR by TR see Figures 1 and 2, and for visualizations of interactions see Supplemental Figures 1 and 2. The exploratory post-hoc voxel-wise comparisons using linear mixed effects modeling indicated there were significant deactivations in the ventral attention network and no significant differences in activations in the dorsal attention network in the feedback by engage condition as compared with the no feedback by engage condition (Table S2). In the feedback by disengage condition, there were significant positive activations in regions consistent with the dorsal attention network and ventral attention networks as compared to the no feedback by disengage condition (Table S3). See Supplemental Results for details.

**Figure 1.**
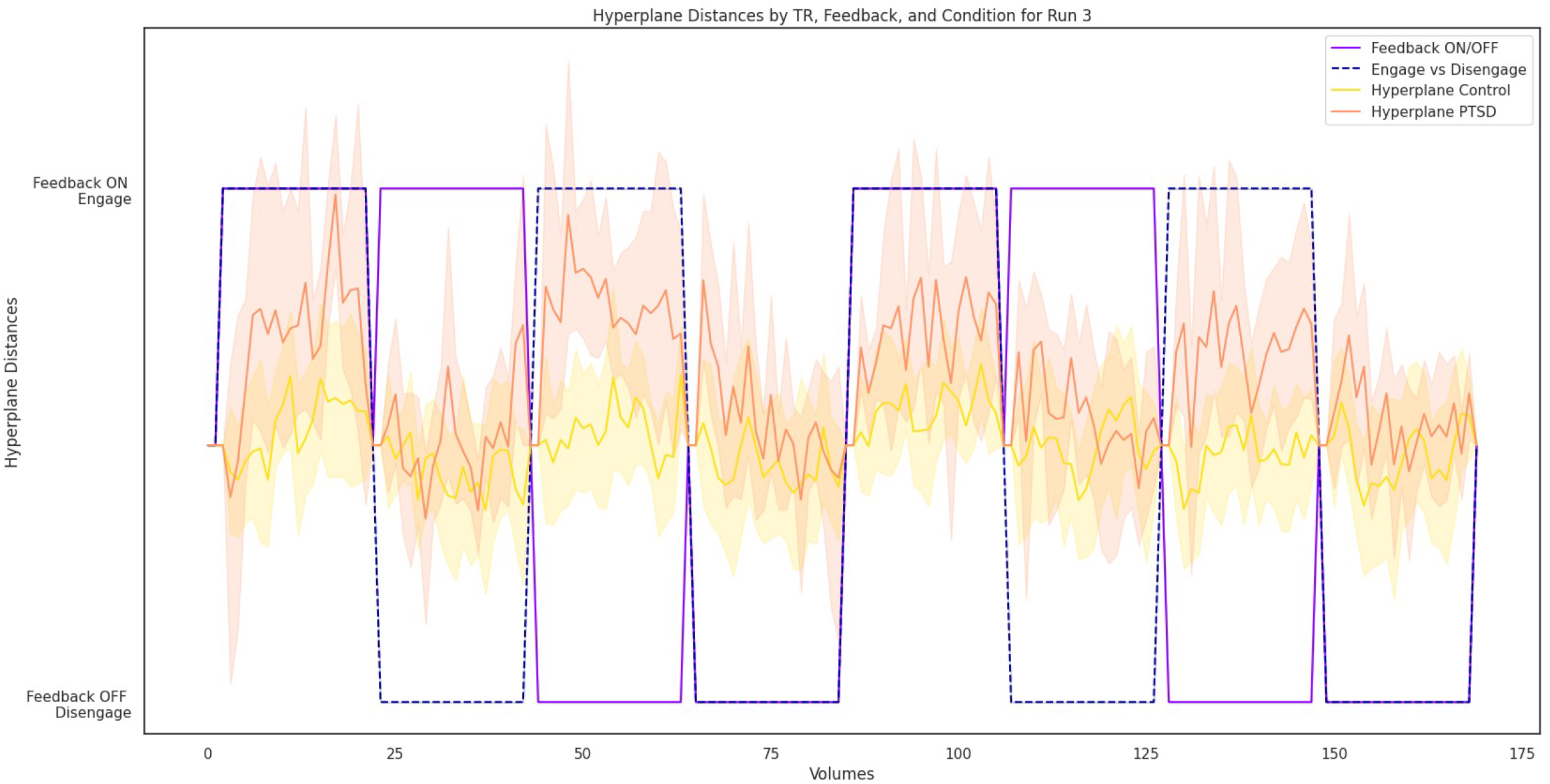
Hyperplane Distance by TR, Feedback, and Condition for Run 3

**Figure 2.**
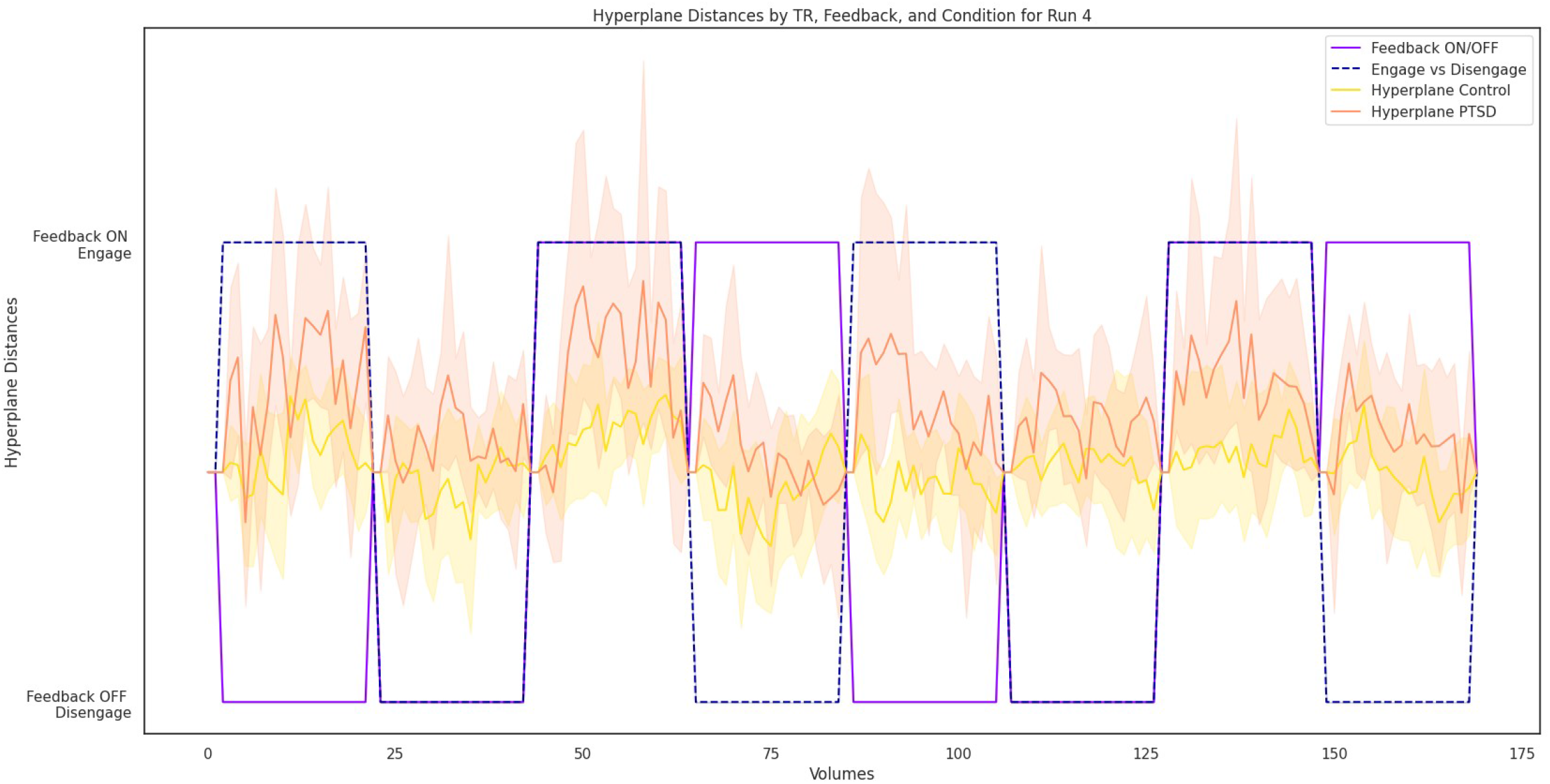
Hyperplane Distance by TR, Feedback, and Condition for Run 4

**Table 1.**
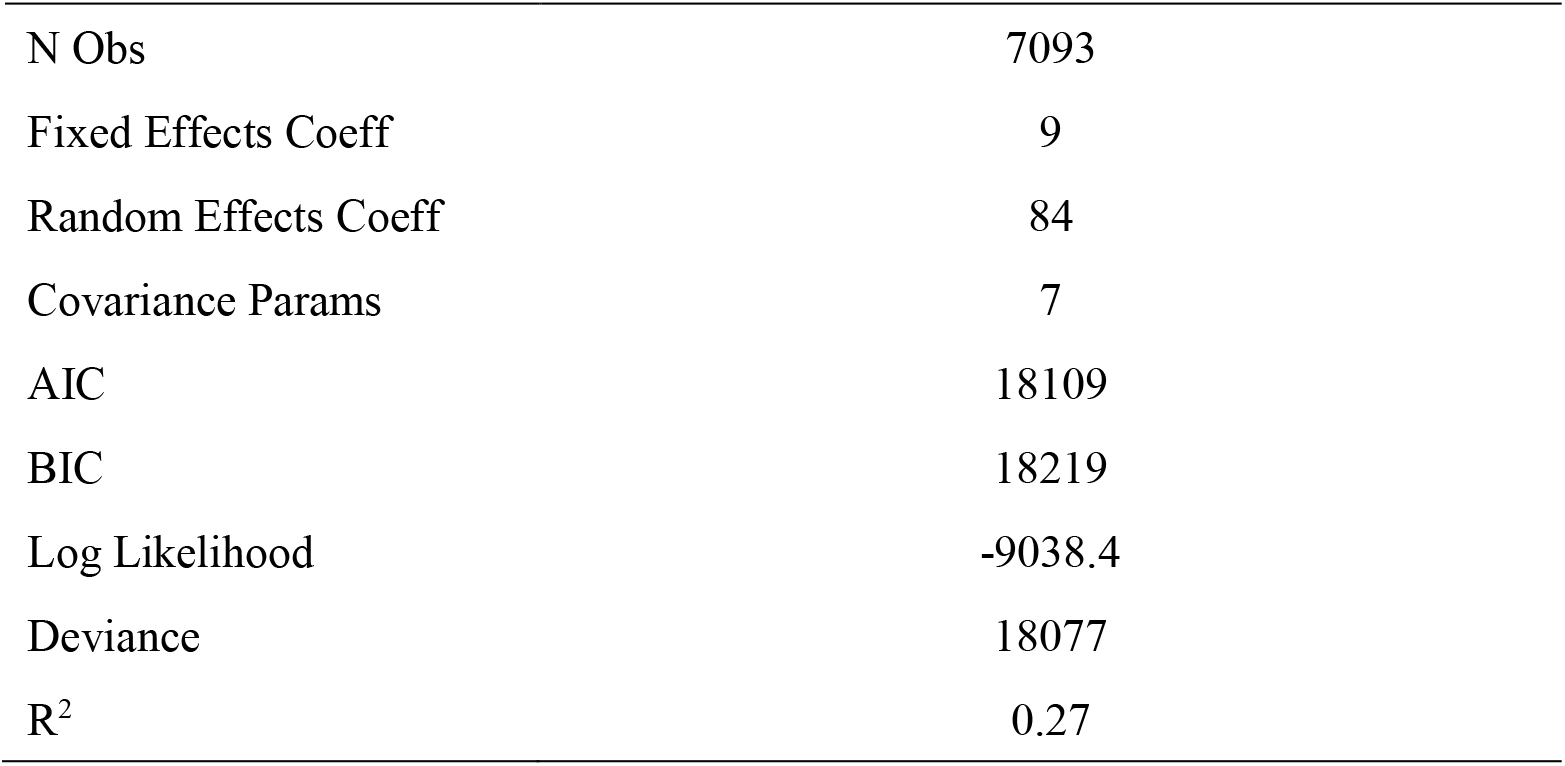
Full Linear Mixed Effects Model Fit Statistics for Hyperplane Distances

**Table 2.** Full Linear Mixed Effects and Simple Slopes Model for Hyperplane Distances

| Full Linear Mixed Effects Model ($n = 21$) | | | | | | | |
| --- | --- | --- | --- | --- | --- | --- | --- |
| Variable | Estimate | SE | $t$ | DF | $p$ | Lower | Upper |
| (Intercept) | -0.30 | 0.12 | -2.51 | 7084 | 0.01 | -0.54 | -0.07 |
| CDN<sup>a</sup> | 0.08 | 0.13 | 0.61 | 7084 | 0.54 | -0.18 | 0.34 |
| FB<sup>b</sup> | -0.05 | 0.05 | -1.03 | 7084 | 0.30 | -0.14 | 0.04 |
| RUN<sup>c</sup> | 0.03 | 0.02 | 1.26 | 7084 | 0.21 | -0.01 | 0.07 |
| GRP<sup>d</sup> | 0.29 | 0.17 | 1.68 | 7084 | 0.09 | -0.05 | 0.63 |
| CDN: FB | 0.22 | 0.05 | 3.99 | 7084 | 0.00 | 0.11 | 0.32 |
| CDN: GRP | 0.38 | 0.19 | 2.01 | 7084 | 0.04 | 0.01 | 0.76 |
| FB: GRP | 0.01 | 0.09 | 0.06 | 7084 | 0.95 | -0.18 | 0.19 |
| CDN: FB: GRP | -0.10 | 0.07 | -1.30 | 7084 | 0.19 | -0.24 | 0.05 |
| cov<sup>e</sup> SUB:CDN | -0.29 | 0.41 |  |  |  | 0.30 | 0.57 |
| cov SUB:FB | 0.65 | 0.17 |  |  |  | 0.11 | 0.25 |
| cov error | 0.85 |  |  |  |  | 0.84 | 0.86 |

| Simple Effect Models |  |  |  |  |  |  |
| --- | --- | --- | --- | --- | --- | --- |
| | Estimate | SE | $t^*$ | $p$ | Lower | Upper |
| Condition: Engage (*no. observations* = 3360) |  |  |  |  |  |  |
| (Intercept) | 0.20 | 0.07 | 2.81 | 0.01 | 0.06 | 0.33 |
| FB | 0.07 | 0.02 | 3.91 | 0.00 | 0.04 | 0.10 |
| Condition: Disengage (*no. observations* = 3733) |  |  |  |  |  |  |
| (Intercept) | 0.04 | 0.05 | 0.80 | 0.43 | -0.06 | 0.13 |
| FB | -0.02 | 0.01 | -1.27 | 0.20 | -0.05 | 0.01 |
| Simple Slopes Difference (*no. observations* = 7093) |  |  |  |  |  |  |
|  | .09 | 0.01 | 6.11 | 0.00 |  |  |
| Condition: Engage (*no. observations* = 3360) |  |  |  |  |  |  |
| (Intercept) | 0.06 | 0.08 | 0.82 | 0.41 | -0.09 | 0.22 |
| GRP | 0.35 | 0.11 | 2.95 | 0.01 | 0.12 | 0.57 |
| Condition: Disengage (*no. observations* = 3733) |  |  |  |  |  |  |
| (Intercept) | -0.04 | 0.07 | -0.56 | 0.58 | -0.16 | 0.09 |
| GRP | 0.14 | 0.09 | 1.50 | 0.14 | -0.04 | 0.32 |
| Simple Slopes Difference (*no. observations* = 7093) |  |  |  |  |  |  |
|  | 0.21 | 0.10 | 2.01 | 0.02 |  |  |
| Group: Control (*no. observations* = 3713) |  |  |  |  |  |  |
| (Intercept) | -0.04 | 0.06 | -0.59 | 0.55 | -0.16 | 0.08 |
| CND | 0.10 | 0.01 | 7.96 | 0.00 | 0.08 | 0.13 |
| Group: PTSD (*no. observations* = 3380) |  |  |  |  |  |  |
| (Intercept) | 0.10 | 0.07 | 1.51 | 0.13 | -0.03 | 0.24 |
| CND | 0.31 | 0.02 | 15.56 | 0.00 | 0.27 | 0.35 |
| Simple Slopes Difference (*no. observations* = 7093) |  |  |  |  |  |  |
|  | -0.21 | 0.01 | -2.91 | 0.00 |  |  |

*Note:* <sup>a</sup>CDN: arouse or suppress conditions, <sup>b</sup>FB: neurofeedback on or off, <sup>c</sup>RUN: run3 or run4, <sup>d</sup>GRP: control or PTSD group <sup>e</sup>cov: covariance, \*the mixed linear model uses z; the comparison between models uses t-values

### Objective Convergent Measurement of Arousal in Response to Neurofeedback

Again, a mixed effects linear model (see Tables 3 and 4) was utilized to assess the effect of CDN, FB, RUN, and GRP on arousal as measured by skin conductance response (SCR). In this model, the main effect of RUN (fixed effect: r = −0.006, *p* = .002, Wald *t*, null: β = 0) was significant where participants had the highest skin conductance response during the third run. There were significant interaction effects: CDN x FB (fixed effect: r = 0.029, *p* = 0.000, Wald *t*, null: β = 0) and CDN x FB x GRP (fixed effect: r = −0.032, *p* < 0.0003, Wald *t*, null: β = 0). Simple effect models revealed feedback was positively associated with skin conductance response in the engage condition (fixed effect: r = 0.02, *p* = 0.0000, z *=* 4.15, null: β = 0), but the difference between SCR in the engage and disengage condition was significant [*t* (6,705) = 2.85, *p* = 0.01]. Simple effects also indicated there were no differences among the groups in the engage (fixed effect: r = −0.01, *p* = 0.56, z *=* −0.58, null: β = 0) or disengage conditions (fixed effect: r = 0.01, *p* = 0.41, z = 0.82, null: β = 0), but the differences between the two effects were significant [*t* (6,705) = −1.99, *p* = 0.02]. The control group’s SCR was higher in the engage condition versus the disengage condition (fixed effect: r = 0.02, *p* = 0.00, *t* = 6.55, null: β = 0) and was higher in the feedback ON condition versus the feedback OFF condition (fixed effect: r = 0.01, *p* = 0.00, z *=* 3.96, null: β = 0). The PTSD group’s SCR followed this same pattern (engage condition fixed effect: r = 0.01, *p* = 0.01, z *=* 2.72, null: β = 0; feedback ON condition fixed effect: r = 0.01, *p* = 0.00, *t* = 3.06, null: β = 0)]. There were no significant differences between SCR in the control or PTSD groups for either feedback ON or engage conditions. The full mixed effects linear model accounted for 5% of the variance in SCR equating to a small effect size (*R^2^* = 0.05). For all simple effect comparisons see Table 4, and for visualization of interactions among variables see Supplemental Figures 3, 4, and 5.

**Table 3.** Full Linear Mixed Effects Model Fit Statistics for SCR

| Table 3. Full Linear Mixed Effects Model Fit Statistics for SCR |  |
| --- | --- |
| N Obs | 6705 |
| Fixed Effects Coeff | 9 |
| Random Effects Coeff | 84 |
| Covariance Params | 7 |
| AIC | -12905 |
| BIC | -12796 |
| Log Likelihood | 6468.5 |
| Deviance | -12937 |
| R<sup>2</sup> | 0.05 |

**Table 4.**
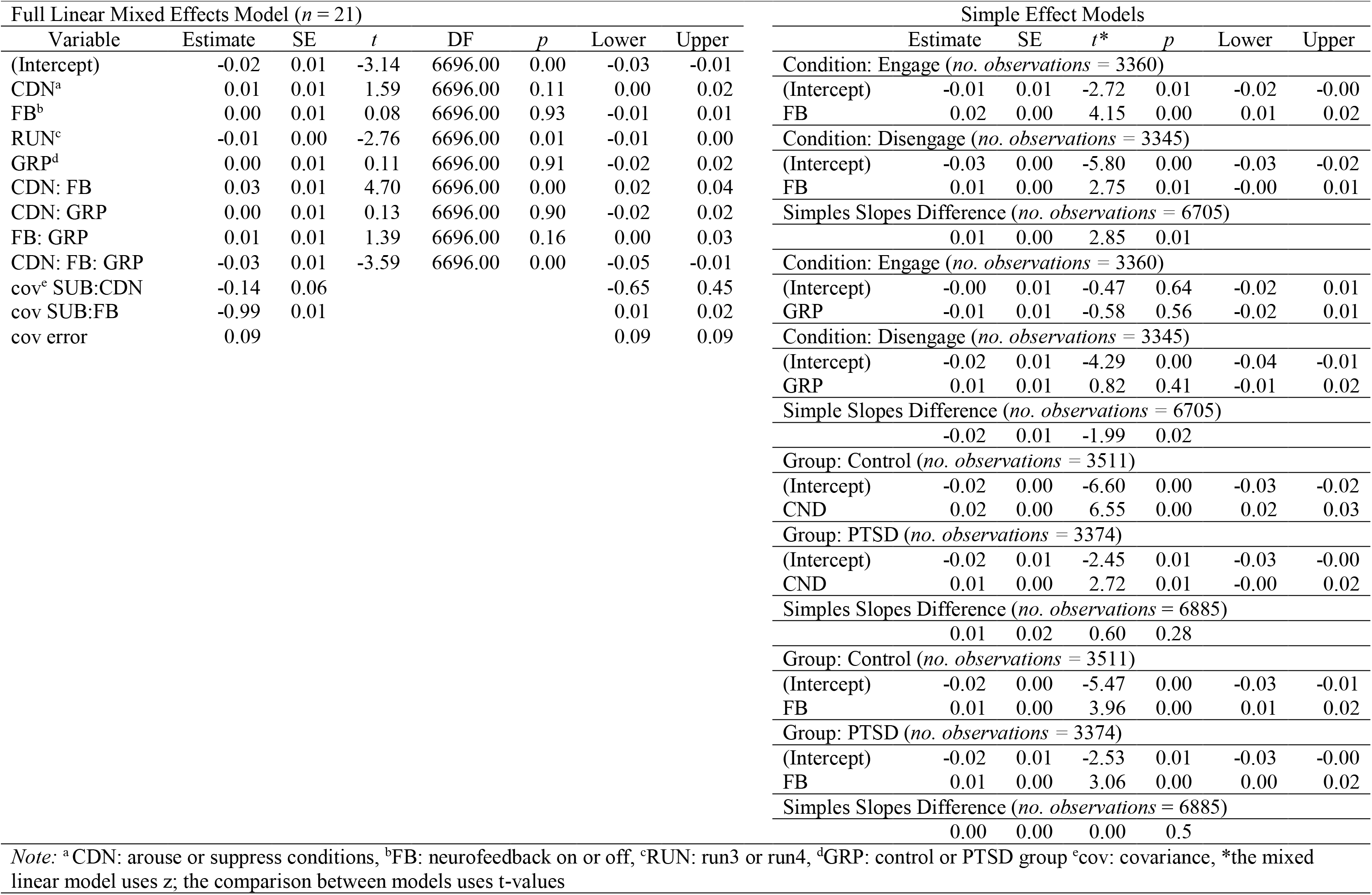
Full Linear Mixed Effects and Simple Slopes Model for SCR

| Full Linear Mixed Effects Model ($n = 21$) | | | | | | | |
| --- | --- | --- | --- | --- | --- | --- | --- |
| Variable | Estimate | SE | $t$ | DF | $p$ | Lower | Upper |
| (Intercept) | -0.02 | 0.01 | -3.14 | 6696.00 | 0.00 | -0.03 | -0.01 |
| CDN<sup>a</sup> | 0.01 | 0.01 | 1.59 | 6696.00 | 0.11 | 0.00 | 0.02 |
| FB<sup>b</sup> | 0.00 | 0.01 | 0.08 | 6696.00 | 0.93 | -0.01 | 0.01 |
| RUN<sup>c</sup> | -0.01 | 0.00 | -2.76 | 6696.00 | 0.01 | -0.01 | 0.00 |
| GRP<sup>d</sup> | 0.00 | 0.01 | 0.11 | 6696.00 | 0.91 | -0.02 | 0.02 |
| CDN: FB | 0.03 | 0.01 | 4.70 | 6696.00 | 0.00 | 0.02 | 0.04 |
| CDN: GRP | 0.00 | 0.01 | 0.13 | 6696.00 | 0.90 | -0.02 | 0.02 |
| FB: GRP | 0.01 | 0.01 | 1.39 | 6696.00 | 0.16 | 0.00 | 0.03 |
| CDN: FB: GRP | -0.03 | 0.01 | -3.59 | 6696.00 | 0.00 | -0.05 | -0.01 |
| cov<sup>e</sup> SUB:CDN | -0.14 | 0.06 |  |  |  | -0.65 | 0.45 |
| cov SUB:FB | -0.99 | 0.01 |  |  |  | 0.01 | 0.02 |
| cov error | 0.09 |  |  |  |  | 0.09 | 0.09 |

| Simple Effect Models |  |  |  |  |  |  |
| --- | --- | --- | --- | --- | --- | --- |
| | Estimate | SE | $t^*$ | $p$ | Lower | Upper |
| Condition: Engage (*no. observations* = 3360) |  |  |  |  |  |  |
| (Intercept) | -0.01 | 0.01 | -2.72 | 0.01 | -0.02 | -0.00 |
| FB | 0.02 | 0.00 | 4.15 | 0.00 | 0.01 | 0.02 |
| Condition: Disengage (*no. observations* = 3345) |  |  |  |  |  |  |
| (Intercept) | -0.03 | 0.00 | -5.80 | 0.00 | -0.03 | -0.02 |
| FB | 0.01 | 0.00 | 2.75 | 0.01 | -0.00 | 0.01 |
| Simple Slopes Difference (*no. observations* = 6705) |  |  |  |  |  |  |
|  | 0.01 | 0.00 | 2.85 | 0.01 |  |  |
| Condition: Engage (*no. observations* = 3360) |  |  |  |  |  |  |
| (Intercept) | -0.00 | 0.01 | -0.47 | 0.64 | -0.02 | 0.01 |
| GRP | -0.01 | 0.01 | -0.58 | 0.56 | -0.02 | 0.01 |
| Condition: Disengage (*no. observations* = 3345) |  |  |  |  |  |  |
| (Intercept) | -0.02 | 0.01 | -4.29 | 0.00 | -0.04 | -0.01 |
| GRP | 0.01 | 0.01 | 0.82 | 0.41 | -0.01 | 0.02 |
| Simple Slopes Difference (*no. observations* = 6705) |  |  |  |  |  |  |
|  | -0.02 | 0.01 | -1.99 | 0.02 |  |  |
| Group: Control (*no. observations* = 3511) |  |  |  |  |  |  |
| (Intercept) | -0.02 | 0.00 | -6.60 | 0.00 | -0.03 | -0.02 |
| CND | 0.02 | 0.00 | 6.55 | 0.00 | 0.02 | 0.03 |
| Group: PTSD (*no. observations* = 3374) |  |  |  |  |  |  |
| (Intercept) | -0.02 | 0.01 | -2.45 | 0.01 | -0.03 | -0.00 |
| CND | 0.01 | 0.00 | 2.72 | 0.01 | -0.00 | 0.02 |
| Simple Slopes Difference (*no. observations* = 6885) |  |  |  |  |  |  |
|  | 0.01 | 0.02 | 0.60 | 0.28 |  |  |
| Group: Control (*no. observations* = 3511) |  |  |  |  |  |  |
| (Intercept) | -0.02 | 0.00 | -5.47 | 0.00 | -0.03 | -0.01 |
| FB | 0.01 | 0.00 | 3.96 | 0.00 | 0.01 | 0.02 |
| Group: PTSD (*no. observations* = 3374) |  |  |  |  |  |  |
| (Intercept) | -0.02 | 0.01 | -2.53 | 0.01 | -0.03 | -0.00 |
| FB | 0.01 | 0.00 | 3.06 | 0.00 | 0.00 | 0.02 |
| Simple Slopes Difference (*no. observations* = 6885) |  |  |  |  |  |  |
|  | 0.00 | 0.00 | 0.00 | 0.5 |  |  |

Note: <sup>a</sup>CDN: arouse or suppress conditions, <sup>b</sup>FB: neurofeedback on or off, <sup>c</sup>RUN: run3 or run4, <sup>d</sup>GRP: control or PTSD group <sup>e</sup>cov: covariance, \*the mixed linear model uses z; the comparison between models uses t-values

## Discussion

There are moderate rates of PTSD remission and improved quality of life after treatment with gold-standard exposure-based treatments ^8,15^ These findings suggest that clinical scientists need to further understand the mediating neural mechanisms of PTSD in order to improve treatment outcomes. Literature contributing to this knowledge base is accumulating, but it largely focuses on regulation of amygdala activity. The current study is the first of which we are aware to utilize a whole brain symptom provocation rt-fMRI-nf paradigm. Results of this study indicate that participants can utilize whole brain-state feedback to engage and disengage with their trauma memory as directed.

In this study we used supervised machine learning to decode whole brain activity during engagement and disengagement with the participant’s trauma memory. We then utilized hyperplane distances to provide participants feedback on their level of engagement or disengagement with the memory. In the mixed effects linear model, there were no main effects for condition, group, or neurofeedback; however, there were significant interaction effects. First, the group by condition interaction was significant and follow-up analyses suggested that both the control and PTSD groups had greater hyperplane distances in the engage condition than in the disengage condition indicating participants evidenced greater engagement with the stimuli in the engage condition than the disengage condition. Moreover, the PTSD group had significantly greater, positive hyperplane distances in the engage condition than the control group. This may suggest that the paradigm was successful at inducing trauma symptoms like hyperarousal and re-experiencing in participants with PTSD^41–43^. It may also indicate that the trauma memory is more engaging than a stress memory.

Additionally, the condition by feedback interaction was also significant. Follow-up analyses indicated that the engage with feedback condition had positive hyperplane distances while the disengage with feedback condition had negative hyperplane distances, and the difference between these effects was significant. However, only the engage condition was significantly associated with participants’ hyperplane distances. This indicates that participants were more engaged with the script in the engage condition while receiving feedback as compared with the engage condition without feedback, and that feedback may not have helped participants engage less in the disengage condition. These results may indicate that whole-brain neurofeedback is effective at increasing engagement with a stressful memory, but not effective at decreasing engagement with that same memory. The null finding during the disengage condition is consistent with efficacy literature on relaxation; specifically, that relaxation is more effective with increased practice and therapeutic sessions.^55^ Overall, our findings largely support the efficacy of rt-fMRI-nf ^28–35^.

Because we utilized a whole-brain multivariate pattern classifier approach to rt-fMRI-nf, rather than a region-specific approach rt-fMRI-nf, we wanted to determine if the significant effects in the feedback by engage versus disengage conditions were related to the use of attentional resources. In the feedback condition while engaging with the trauma/stress memory, participants displayed deactivations in the ventral attention network as compared with the no feedback condition while engaging with the memory. This may suggest that when individuals do not receive feedback while engaging with the trauma/stress memory they may need to reorient to the task more often than when they receive feedback.^56^ In the feedback condition while disengaging from the trauma/stress memory, participants displayed more positive activations in the dorsal and ventral attention networks than in the no feedback condition while disengaging with the memory. This may suggest that feedback while trying to disengage from the memory utilizes more top down regulation and reorienting^56^ than no feedback while trying to disengage from the memory. Please see Supplemental materials for specific analysis and results details.

To independently test the effect of rt-fMRI-nf on a convergent measure of physiological arousal, we utilized participant’s skin conductance responses. Again, there were no significant main effects for feedback, condition, or group, but there was a significant main effect of run where participants evidenced greater skin conductance response (SCR) in the earlier run than the later run. This is particularly important in the context of PTSD treatment. In exposure-based treatments, the individual reads or describes their trauma experience successively with the goal of extinguishing (or greatly reducing) the fear associated with that memory (i.e., extinction^12,58,59^). Because arousal decreased after repeated presentations of the participant’s trauma/stress script, it is possible that extinction occurred in our participants which would be consistent with the experimental fear extinction literature indicating skin conductance responses decrease in healthy controls, trauma-exposed controls, and participants with PTSD during extinction learning ^60–63^. Importantly, the PTSD clinical literature also indicates autonomic arousal declines from pre-to post-treatment after exposure therapy for PTSD ^64^ and that within-session habituation is related to positive treatment outcomes ^65^. Alternatively, decreases in arousal could also be explained by participant fatigue.

In addition to the main effect of run, there was an interaction effect of condition by feedback. Specifically, when participants were provided feedback they had significantly higher arousal during the engage blocks. This result independently supports that rt-fMRI-nf maybe able to increase the effectiveness of exposure therapies through activating trauma-related arousal that would facilitate extinction and exposure to corrective information^12,13^ which are key components to the success of exposure-based therapies ^58^. However, there is a caveat to this finding as the three-way interaction among engagement, feedback, and group suggested that individuals with PTSD evidenced less arousal in the engage condition than the disengage condition, but all participants demonstrated higher arousal in the feedback condition versus the no feedback condition. There were no significant differences between the control and PTSD groups in their levels of arousal during feedback. This three-way interaction could indicate that individuals with PTSD have difficulties disengaging from their trauma memories which is consistent with reexperiencing and hyperarousal symptoms.

Accumulating data indicates that rt-fMRI-nf can be utilized for emotion regulation in healthy individuals ^21^, individuals with symptoms of depression ^23,24^ and anxiety ^25,26^, and there is accumulating evidence that it can also be utilized in patients with PTSD ^27–34^. However, our study extends this literature in many ways. First, because mental states are represented in the brain by distributed activations, our approach utilizing multivariate pattern classification allowed us to decode engagement and disengagement with the stimuli and provide participants feedback that was representative of whole brain patterns of activation ^66^ rather than feedback based on regions of interest. Consequently, the current study is likely more robust as it avoids over reliance on any one ROI ^17^. Secondly, most studies utilize rt-fMRI-nf in male combat veteran populations which may be qualitatively different from women with histories of interpersonal violence ^67,68^, and we demonstrated that rt-fMRI-nf could be utilized in women with interpersonal violence-related PTSD. Finally, we utilized a symptom provocation rt-fMRI-nf paradigm that is most similar to existing gold-standard treatments for PTSD which increases this study’s relevance to increasing the efficacy of existing treatments.

### Limitations

Although we demonstrated typical hemodynamic responses in regions related to symptom provocation studies and rt-fMRI-nf, and we demonstrated that skin conductance response largely reflected arousal with engagement and feedback, we did not measure PTSD symptoms in participants after the study period. Thus, we cannot speak to reductions in PTSD symptoms related to the rt-fMRI-nf paradigm. Our study was a small-n clinical trial, and while the experimental nature of the study increases the rigor, recent studies have indicated that small-n fMRI studies may over estimate effects sizes^69^ which indicate future research is needed to replicate this finding.

### Conclusions

Interpersonal violence is common among women^1^ and it increases the likelihood of developing PTSD when compared to other trauma types^2–4^. Treatment success for PTSD is at roughly 50% which increases the importance of understanding neural mechanisms in order to inform more effective treatments. With the exception of largely consistent activations in the dACC and deactivations in the vmPFC among individuals with PTSD, the literature on neural mechanisms is inconsistent^17^. To aid our understanding of neural mechanisms, rt-fMRI-nf has attempted to modulate regions that are hypothesized to underly PTSD symptoms; however, most studies utilize male combat veterans, paradigms are not tied to current empirically supported treatment mechanisms, and paradigms utilize feedback from regions of interest that may not fully capture the neural mechanisms of PTSD. To fill these gaps in the literature, we utilized a whole-brain multivariate pattern classifier to provide real-time neurofeedback to participants about their level of engagement with their trauma memory. Our findings suggest that even when considering individual variability, participants were better at engaging with their trauma/stress memory when they were given feedback versus when they were not given feedback, and that individuals with PTSD displayed more engagement with the memory than healthy controls. Skin conductance response and brain map activations were used to further validate these findings.

## Supporting information

All Supplemental Materials

## Source Code and Data Availability

The authors have made the bids https://github.com/kabush/rPEP2bids, preprocessing https://github.com/kabush/rPEP_preproc, and analysis code used in for this manuscript publicly available https://github.com/maegancalvert/rPEP. All reported functional neuroanatomical activation maps are publicly available via the study’s Open Science Framework repository: https://osf.io/dkfs2/. For replication purposes only, de-identified raw data from this study may be made privately available upon request.

