## Supplementary material for "Increasing the efficacy of exposure using a whole brain approach to real-time fMRI neurofeedback among female victims of interpersonal violence": All Supplemental Materials

### Increasing the efficacy of exposure using a real-time fMRI neurofeedback paradigm among women with histories of interpersonal violence

Maegan L. Calvert[1], Josh M. Cisler[2], & Keith A. Bush[1]

[1]Department of Psychiatry, Brain Imaging Research Center, University of Arkansas for Medical Sciences

[2]Department of Psychiatry & Behavioral Sciences, Dell Medical School, The University of Texas at Austin

**Supplemental Information**

**Supplemental Methods**

**Supplemental Equation SE1.**

**Supplemental Results**

**Figure S1.** Hyperplane Distances by Disengage versus Engage Separated by Feedback OFF and ON

**Figure S2.** Hyperplane Distances by Disengage versus Engage Separated by Group

**Figure S3.** SCR by Disengage versus Engage Separated by Feedback OFF and ON

**Figure S4.** SCR by Feedback OFF and ON Separated by Group

**Figure S5.** SCR by Disengage versus Engage Separated by Group

**Table S1.** Descriptive Statistics for Clinical Variables by Total Sample and by Group

**Table S2.** Regions of Peak Activation for Feedback ON and OFF in the Engage Condition for Whole-Brain rt-fMRI-nf

**Table S3.** Regions of Peak Activation to Feedback ON and OFF in the Disengage Condition for Whole-Brain rt-fMRI-nf

**Other** CRED-nf Best Practices Checklist

**Supplemental References**

#### Supplemental Methods

##### *Post-hoc Minimal Image Preprocessing with fmriprep*

Results included in this manuscript come from preprocessing performed using FMRIPREP version stable [1,2], a Nipype [3,4] based tool. Each T1w (T1-weighted) volume was corrected for INU (intensity non-uniformity) using N4BiasFieldCorrection v2.1.0 [5] and skull-stripped using antsBrainExtraction.sh v2.1.0 (using the OASIS template). Brain surfaces were reconstructed using recon-all from FreeSurfer v6.0.1 [6], and the brain mask estimated previously was refined with a custom variation of the method to reconcile ANTs-derived and FreeSurfer-derived segmentations of the cortical gray-matter of Mindboggle [7]. Spatial normalization to the ICBM 152 Nonlinear Asymmetrical template version 2009c [8] was performed through nonlinear registration with the antsRegistration tool of ANTs v2.1.0 [9], using brain-extracted versions of both T1w volume and template. Brain tissue segmentation of cerebrospinal fluid (CSF), white-matter (WM) and gray-matter (GM) was performed on the brain-extracted T1w using fast [10].

Functional data was slice time corrected using 3dTshift from AFNI v16.2.07 [11] and motion corrected using mcflirt (FSL v5.0.9 [12]). This was followed by co-registration to the corresponding T1w using boundary-based registration [13] with six degrees of freedom, using bbregister (FreeSurfer v6.0.1). Motion correcting transformations, BOLD-to-T1w transformation and T1w-to-template (MNI) warp were concatenated and applied in a single step using antsApplyTransforms (ANTs v2.1.0) using Lanczos interpolation.

Physiological noise regressors were extracted applying CompCor [14]. Principal components were estimated for the two CompCor variants: temporal (tCompCor) and anatomical (aCompCor). A mask to exclude signal with cortical origin was obtained by eroding the brain mask, ensuring it only contained subcortical structures. Six tCompCor components were then calculated including only the top 5% variable voxels within that subcortical mask. For aCompCor, six components were calculated within the intersection of the subcortical mask and the union of CSF and WM masks calculated in T1w space, after their projection to the native space of each functional run. Frame-wise displacement [15] was calculated for each functional run using the implementation of Nipype. Many internal operations of FMRIPREP use Nilearn [7], principally within the BOLD-processing workflow. For more details of the pipeline see https://fmriprep.readthedocs.io/en/stable/workflows.html.

***Additional Post-hoc MR Image Processing***

Using fmriprep's motion parameter outputs, we completed preprocessing of the fMRI data using AFNI [11] according to the following steps: (1) regression of the mean time courses and temporal derivatives of the white matter and cerebrospinal fluid masks as well as a 24-parameter motion mode (Power et al., 2012; Power et al., 2014), (2) spatial smoothing (8 mm FWHM), (3) detrending, (4) temporal filtering (.0078 Hz high-pass), and (5) scaling to percent signal change.

***Post-hoc Identification of the Neural Correlates of Neurofeedback Processing in the Engage versus Disengage Conditions***

We looked for potential alternate cognitive processes (i.e., attentional processes), concurrent with neurofeedback processing, that may explain our observed FB X CND effect. To do so, we constructed statistical maps of the neural activation patterns that distinguish neurofeedback-guided from unguided engagement/disengagement of the trauma/stress script. To do so we constructed a whole-brain gray matter linear mixed-effects model of neurofeedback processing. We modeled the activation within each GM voxel, i, of each volume, j, and subject, k, as a function of the neurofeedback guidance provided during that volume {on, off} and engage {on, off] as well as relative temporal position of that volume within the fMRI acquisition {scan #3, scan #4} while controlling for random slope and intercept effects subject-wise as shown in Supplemental Equation, SE1, using Wilkinson-Rogers syntax [16] extended for LME [17].

activation[i,j,k] = 1 + neurofeedback[j] + engage[j] + (neurfeedback[j]:engage[j]) + (1| subj[k]) **[SE1]**

We applied AFNI's 3dlme function to solve this equation and outputs both general linear test results of our desired contrast (1*on – 1*off) as well as model residuals. We then computed cluster-size thresholds using previously reported steps [18]: (1) estimate the shape parameters of the of the residuals using the spatial autocorrelation function (ACF) which is built-in to AFNI's 3dFWHMx function, (2) simulate, via 3dClustSim (-acf option), the cluster-size threshold for the statistical map (thresholded to $p<0.001$, uncorrected) that survive family-wise error (FWE) corrected thresholds of $p<0.05$, assuming that clusters were formed from contiguous (face-touching, NN = 1) voxels.

**Supplemental Results**

***Neural Correlates of Neurofeedback Processing in the Engage Condition***

Cluster threshold based on AFNI's 3dClustSim was set at 15 based on the median of model residuals. During feedback ON blocks while engaging with the trauma or stress script, there were only clusters of deactivations in the ventral attention networks relative to the feedback OFF blocks and there were no significant activations or deactivations in the dorsal attention network during feedback ON versus feedback OFF blocks while engaging with the memory. Specifically, the bilateral intraparietal sulcus and the bilateral frontal eye fields from the dorsal attention network[19] did not show significantly different activity in the feedback ON versus OFF condition while engaging with the memory. The superior frontal gyrus within the ventral attention network also did not evidence significantly different activity in the feedback ON versus feedback OFF conditions while engaging with the trauma memory. However, other regions in the ventral attention network like the medial frontal gyrus (IFJ, A9/46v – bilateral), inferior frontal gyrus (A44d, IFS - bilateral), and temporoparietal junction (A40c, A39rv – bilateral) showed significantly less activation in the feedback ON versus feedback OFF conditions while engaging with the trauma/stress memory. This may indicate that the ventral attention network is more active in the feedback OFF versus feedback ON condition while engaging with stressful memories. See Table S2. For activation maps, see OSF repository.

***Neural Correlates of Neurofeedback Processing in the Disengage Condition***

Cluster threshold based on AFNI's 3dClustSim was set at 15 based on model residuals. While disengaging with the trauma/stress memory, there were clusters of positive activations in the ventral and dorsal attention networks in the feedback ON relative to the feedback OFF conditions. The dorsal attention network regions including the left intraparietal sulcus and bilateral frontal eye fields did not show significant differences in the feedback ON versus feedback OFF conditions while disengaging with the trauma/stress memory. However, the right intraparietal sulcus did evidence significant positive activations in the feedback ON versus the feedback OFF conditions while disengaging. In the ventral attention network, the bilateral medial frontal gyrus (A9/46v – left; A9/46v, A8vl, A6vl - right), the bilateral superior frontal gyrus (A6m, A10m – left; A8dl, A9l - right), left inferior frontal gyrus (IFS), and the bilateral temporoparietal junction (A40c, A39rv) displayed significant positive activations in the feedback ON versus feedback OFF conditions while disengaging from the trauma/stress memory. There were no significantly different activations in the right inferior frontal gyrus between the two conditions. This may indicate that during the feedback ON versus feedback OFF conditions while disengaging from the trauma/stress memory, both the dorsal and ventral attention networks are active in attending to feedback and modifying behavior. See Table S3. For activation maps, see OSF repository.

**Figure S1.**

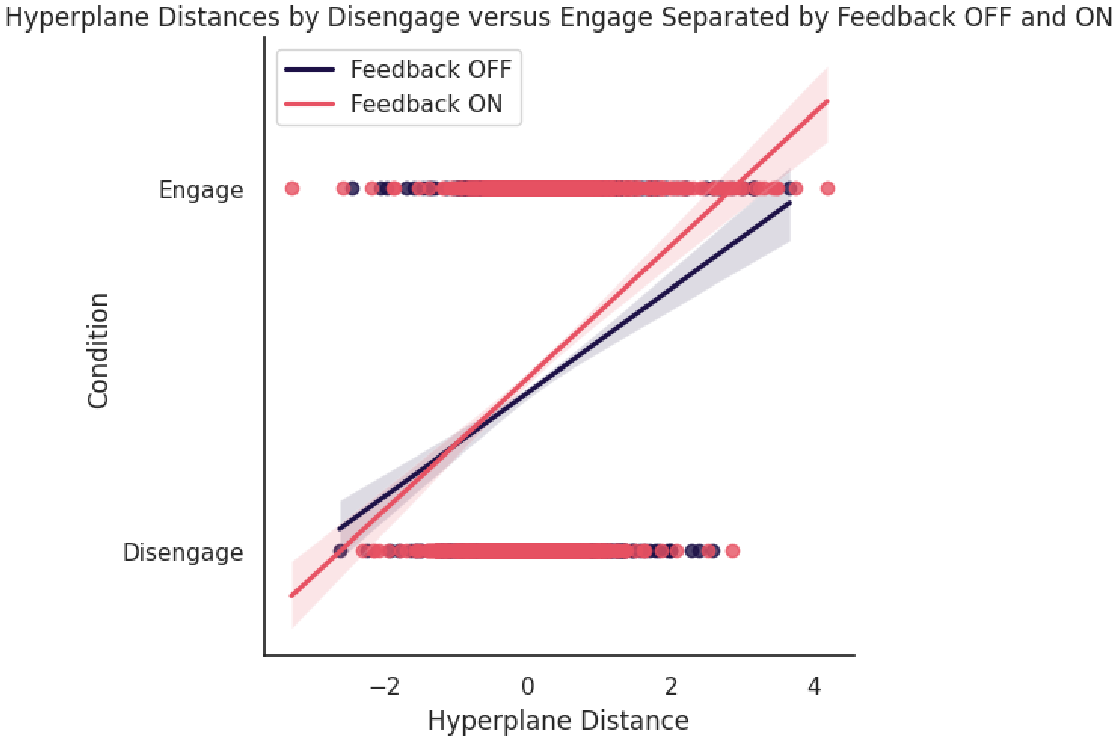


**Figure S2.**

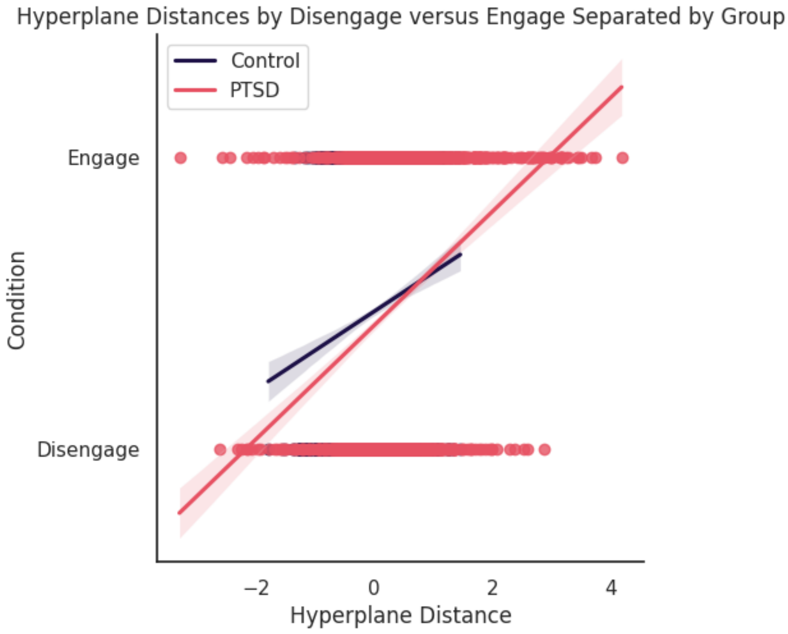

**Figure S3.**

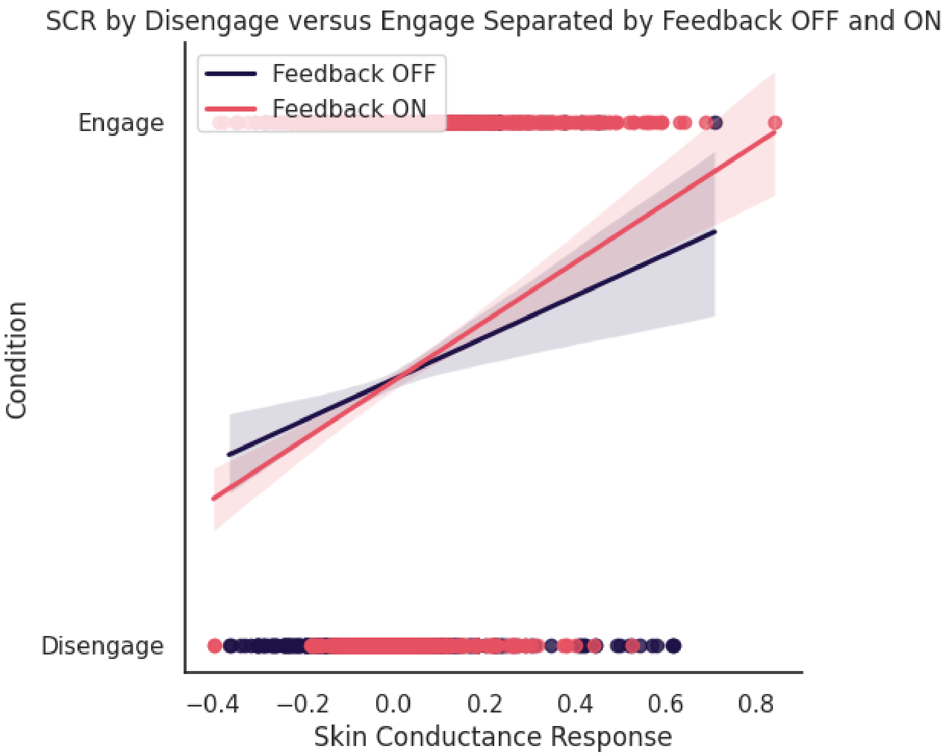


**Figure S4.**

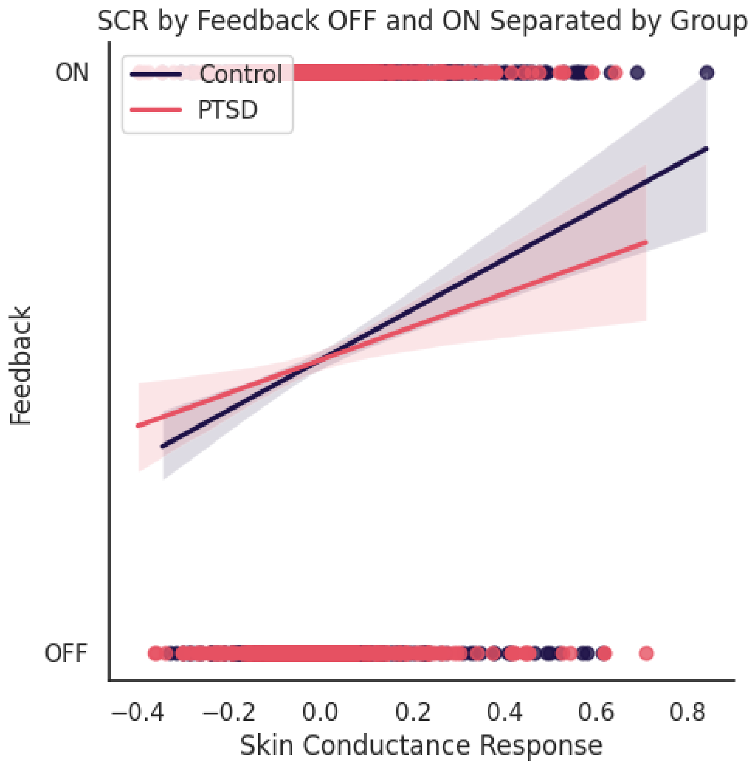

**Figure S5.**

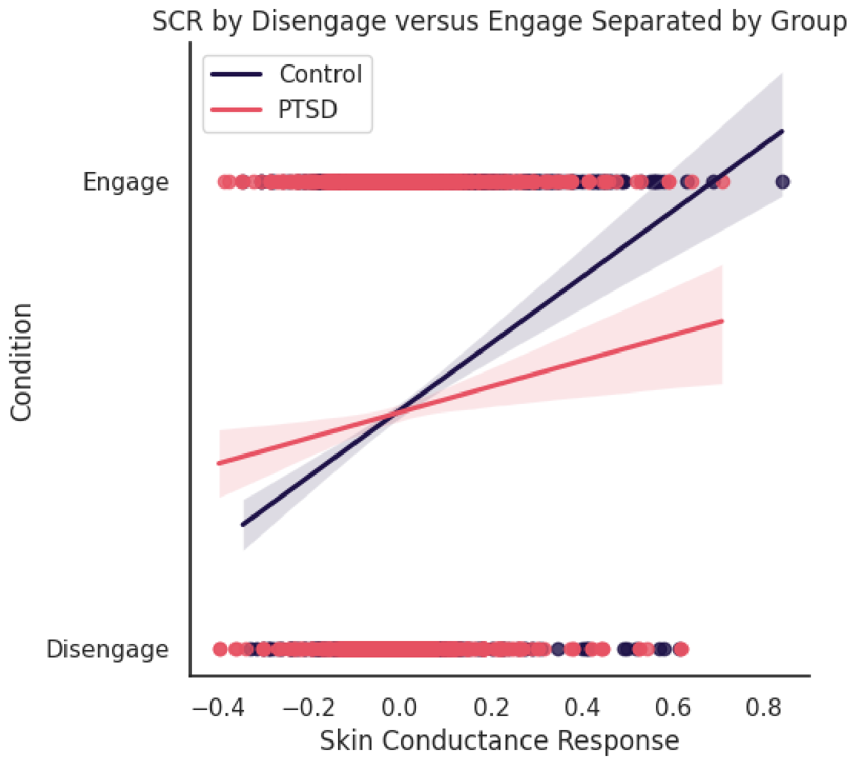
SCR by Disengage versus Engage Separated by Group
Control
PTSD
Engage
Disengage
Condition
−0.4
−0.2
0.0
0.2
0.4
0.6
0.8
Skin Conductance Response

**Table S1. Descriptive Statistics for Demographic and Clinical Variables by Total Sample and by Group**

| Variable | Total *(N)* | Total Mean *(SD)* | Total Min - Max | Control *(n)* | Control Mean *(SD)* | Control Min - Max | PTSD *(n)* | PTSD Mean *(SD)* | PTSD Min - Max | *t* Statistic |
|---|---|---|---|---|---|---|---|---|---|---|
| Age | 21.00 | 32.95 (9.33) | 21.00 - 50.00 | 11.00 | 30.09 (8.49) | 21.00 - 50.00 | 10.00 | 36.10 (9.61) | 21.00 - 49.00 | -1.52 |
| PCL score | 21.00 | 28.71 (22.28) | 0.00 - 67.00 | 11.00 | 10.64 (11.16) | 0.00 - 28.00 | 10.00 | 48.60 (11.24) | 31.00 - 67.00 | -7.76*** |
| CTQ score | 21.00 | 68.29 (17.49) | 32.00 - 101.00 | 11.00 | 64.18 (15.89) | 36.00 - 84.00 | 10.00 | 72.80 (18.87) | 32.00 - 101.00 | -1.14 |
| Any lifetime or current diagnosis | 16.00 | | | 6.00 | | | 10.00 | | | -2.89* |
| Handedness | | | | | | | | | | |
| Left | 2 | | | 2 | | | 0 | | | |
| Right | 17 | | | 9 | | | 8 | | | |
| Race | | | | | | | | | | |
| White | 16 | | | 9 | | | 7 | | | |
| Black | 3 | | | 2 | | | 1 | | | |
| Latina | 1 | | | 0 | | | 1 | | | |
| Asian American | 1 | | | 0 | | | 1 | | | |
| Education | | | | | | | | | | |
| <12 years | 2 | | | 0 | | | 2 | | | |
| >12 years | 19 | | | 11 | | | 8 | | | |
| Poly-victimization | 16 | | | 8 | | | 8 | | | |
| Treatment | | | | | | | | | | |
| Medications | 4 | | | 0 | | | 4 | | | |
| Therapy | 6 | | | 2 | | | 4 | | | |

*Note: * p < .05 *** p< .001*

**Table S2. Regions of Peak Activation for Feedback ON and OFF in the Engage Condition for Whole-Brain rt-fMRI-nf**

| Cluster Number | AAL Label | BA# | Human Braintomme (% overlap) | | Voxels | *x* | *y* | *z* |
|---|---|---|---|---|---|---|---|---|
| 1 | Temporal_Mid_R | 37 | A37mv_right | 20.80% | 361 | 60 | -55 | 3 |
| | | | A37dl_right | 19.60% | | | | |
| 2 | Cerebelum_6_L | 18 | A37mv_left | 36.20% | 287 | -25 | -76 | -19 |
| | | | A37lv_left | 29.00% | | | | |
| 3 | Precentral_R | 6 | A6cvl_right | 43.50% | 134 | 45 | 6 | 33 |
| | | | A44d_right | 31.10% | | | | |
| 4 | Frontal_Inf_Tri_R | 46 | A9/46v_right | 55.80% | 130 | 48 | 45 | -1 |
| | | | IFS_right | 26.10% | | | | |
| 5 | Occipital_Mid_L | 18 | mOccG_left | 47.20% | 97 | -31 | -88 | 6 |
| | | | iOccG_left | 36.50% | | | | |
| 6 | Occipital_Mid_R | 18 | mOccG_right | 76.50% | 87 | 27 | -97 | 15 |
| | | | OPC_right | 22.90% | | | | |
| 7 | Parietal_Inf_R | 7 | A40rd_right | 45.30% | 73 | 39 | -46 | 45 |
| | | | A7ip_right | 27.60% | | | | |
| 8 | Angular_R | 39 | A40c_right | 99.00% | 61 | 63 | -49 | 36 |
| | | | A40rd_right | 0.40% | | | | |
| **9** | **Cuneus_L** | **31** | A31_left | 43.20% | **57** | **-4** | **-67** | **27** |
| | | | A31_right | 22.90% | | | | |
| 10 | Precentral_L | 44 | A6cvl_left | 57.60% | 54 | -46 | 3 | 24 |
| | | | IFJ_left | 36.60% | | | | |
| 11 | Insula_R | 13 | dIa_right | 39.30% | 53 | 33 | 27 | 3 |
| | | | A44op_right | 30.50% | | | | |
| 12 | Parietal_Inf_L | 39 | A40c_left | 84.90% | 53 | -58 | -46 | 45 |
| | | | A40rd_left | 5.30% | | | | |
| **13** | **Temporal_Sup_L** | **41** | TE1.0_1.2_left | 54.10% | **40** | **-58** | **-7** | **6** |
| | | | A1/2/3tonIa_left | 23.70% | | | | |
| **14** | **Insula_R** | **1** | A1/2/3tonIa_right | 51.20% | **23** | **39** | **-13** | **18** |
| | | | dIg_right | 37.10% | | | | |
| 15 | Frontal_Inf_Tri_L | 46 | A9/46v_left | 93.00% | 21 | -46 | 45 | 15 |
|  |  |  | --- | ----- |  |  |  |  |
| 16 | Frontal_Mid_L | 10 | A46_left | 86.20% | 19 | -31 | 63 | 12 |
| | | | A9/46v_left | 5.10% | | | | |
| 17 | Frontal_Mid_Orb_R | 10 | A46_right | 54.20% | 18 | 33 | 60 | -1 |
| | | | A10l_right | 45.80% | | | | |

*Note:* Bold clusters are positive. BrainSpy was utilized to generate AAL and BA# locations. AFNI whereami was utilized to generate Human Brainnetome locations.

**Table S3. Regions of Peak Activation to Feedback ON and OFF in the Disengage Condition for Whole-Brain rt-fMRI-nf**

| Cluster Number | AAL Label | BA# | Human Braintomme (% overlap) | | Voxels | *x* | *y* | *z* |
|---|---|---|---|---|---|---|---|---|
| 1 | SupraMarginal_R | 40 | A40c_right | 11.20% | 1269 | 66 | -37 | 36 |
| | | | A39rv_right | 9.00% | | | | |
| 2 | Temporal_Mid_L | 39 | A40c_left | 14.40% | 739 | -58 | -64 | 12 |
| | | | A37dl_left | 12.30% | | | | |
| 3 | Occipital_Mid_R | 39 | lsOccG_right | 56.70% | 139 | 33 | -79 | 24 |
| | | | mOccG_right | 21.40% | | | | |
| 4 | SupraMarginal_L | 40 | TE1.0_1.2_left | 37.90% | 122 | -52 | -25 | 18 |
| | | | A40rv_left | 30.10% | | | | |
| 5 | Occipital_Inf_R | 37 | V5/MT+_right | 45.70% | 92 | 48 | -67 | -13 |
| | | | A37lv_right | 23.70% | | | | |
| 6 | Cerebelum_6_R | 37 | A37mv_right | 48.50% | 65 | 36 | -55 | -22 |
| | | | rLinG_right | 13.80% | | | | |
| 7 | Frontal_Mid_R | 10 | A46_right | 40.80% | 56 | 39 | 54 | 15 |
| | | | A9/46v_right | 40.70% | | | | |
| 8 | Postcentral_R | 4 | A4hf_right | 60.50% | 56 | 54 | -7 | 27 |
| | | | A1/2/3ulhf_right | 38.10% | | | | |
| 9 | Frontal_Inf_Tri_L | 46 | IFS_left | 61.50% | 54 | -52 | 33 | 21 |
| | | | A9/46v_left | 36.00% | | | | |
| 10 | Parietal_Sup_L | 7 | A7pc_left | 49.30% | 54 | -22 | -46 | 72 |
| | | | A7r_left | 23.70% | | | | |
| 11 | Occipital_Mid_R | 19 | mOccG_right | 96.40% | 52 | 36 | -85 | 9 |
| | | | V5/MT+_right | 2.70% | | | | |
| 12 | Precentral_L | 6 | A6cdl_left | 88.10% | 47 | -19 | -13 | 63 |
| | | | A4t_left | 2.00% | | | | |
| 13 | Frontal_Sup_Medial_L | 10 | A10m_left | 80.00% | 41 | -4 | 57 | 12 |
| | | | A32p_left | 20.00% | | | | |
| 14 | Calcarine_L | 17 | vmPOS_left | 42.30% | 40 | -10 | -67 | 9 |
| | | | rCunG_left | 33.30% | | | | |
| 15 | Cerebelum_Crus1_R | 37 | A37mv_right | 51.60% | 37 | 36 | -76 | -19 |
| | | | iOccG_right | 1.80% | | | | |
| 16 | Parietal_Sup_L | 7 | A39rd_left | 58.30% | 37 | -25 | -73 | 60 |
| | | | A7ip_left | 8.00% | | | | |
| 17 | Cerebelum_Crus1_L | 18 | cLinG_left | 0.10% | 36 | -7 | -79 | -19 |
| 18 | Temporal_Mid_R | 37 | A37dl_right | 62.10% | 31 | 57 | -58 | 9 |
| | | | A39c_right | 24.90% | | | | |
| 19 | Frontal_Mid_R | 8 | A8dl_right | 53.30% | 30 | 30 | 21 | 54 |
| | | | A6vl_right | 41.80% | | | | |
| 20 | Occipital_Sup_L | 19 | A7c_left | 52.70% | 29 | -16 | -85 | 45 |
| | | | lsOccG_left | 28.90% | | | | |

| | | | | | | | | |
|---|---|---|---|---|---|---|---|---|
| 21 | Cingulum_Ant_R | 9 | A32sg_right | 54.30% | 27 | 9 | 42 | 18 |
| | | | A10m_right | 22.40% | | | | |
| 22 | Frontal_Sup_R | 8 | A9l_right | 63.30% | 24 | 12 | 36 | 51 |
| | | | A8dl_right | 34.90% | | | | |
| 23 | Putamen_R | 49 | vCa_right | 45.30% | 21 | 21 | 9 | -10 |
| | | | NAC_right | 34.90% | | | | |
| 24 | Frontal_Mid_R | 9 | A8vl_right | 92.60% | 19 | 45 | 30 | 33 |
| | | | A9/46d_right | 6.00% | | | | |
| 25 | Temporal_Pole_Sup_R | 44 | TE1.0_1.2_right | 79.80% | 18 | 54 | 9 | -4 |
| | | | A4tl_right | 11.30% | | | | |
| 26 | Occipital_Mid_L | 39 | A39rv_left | 95.20% | 18 | -43 | -79 | 36 |
| | | | A39c_left | 3.00% | | | | |
| 27 | Temporal_Mid_L | 21 | cpSTS_left | 87.10% | 16 | -46 | -52 | 12 |
| | | | A39rv_left | 12.30% | | | | |
| 28 | Fusiform_L | 19 | A37mv_left | 62.10% | 15 | -25 | -73 | -7 |
| | | | rLinG_left | 31.00% | | | | |
| 29 | Frontal_Sup_Medial_R | 9 | A9m_right | 68.40% | 15 | 6 | 51 | 27 |
| | | | A10m_right | 31.60% | | | | |

*Note:* All clusters are positive. BrainSpy was utilized to generate AAL and BA# locations. AFNI whereami was utilized to generate Human Brainnetome locations.

| CRED-nf best practices checklist 2020 | | | |
|---|---|---|---|
| **Domain** | **Item #** | **Checklist item** | **Reported on page #** |
| **Pre-experiment** | | | |
| | 1a | Pre-register experimental protocol and planned analyses | Pg. 7 |
| | 1b | Justify sample size | |
| **Control groups** | | | |
| | 2a | Employ control group(s) or control condition(s) | Pg. 7 |
| | 2b | When leveraging experimental designs where a double-blind is possible, use a double-blind | N/A |
| | 2c | Blind those who rate the outcomes, and when possible, the statisticians involved | N/A |
| | 2d | Examine to what extent participants and experimenters remain blinded | N/A |
| | 2e | In clinical efficacy studies, employ a standard-of-care intervention group as a benchmark for improvement | N/A |
| **Control measures** | | | |
| | 3a | Collect data on psychosocial factors | Pg. 7/ S1 |
| | 3b | Report whether participants were provided with a strategy | Pg. 8 |
| | 3c | Report the strategies participants used | Pg. 9 |
| | 3d | Report methods used for online-data processing and artefact correction | Pg. 9 |
| | 3e | Report condition and group effects for artefacts | |
| **Feedback specifications** | | | |
| | 4a | Report how the online-feature extraction was defined | Pg. 10 |
| | 4b | Report and justify the reinforcement schedule | |
| | 4c | Report the feedback modality and content | Pg. 8-9 |
| | 4d | Collect and report all brain activity variable(s) and/or contrasts used for feedback, as displayed to experimental participants | Pg. 10 |
| | 4e | Report the hardware and software used | Pg. 8 |
| **Outcome measures** | | | |
| Brain | 5a | Report neurofeedback regulation success based on the feedback signal | Pg. 11 |
| | 5b | Plot within-session and between-session regulation blocks of feedback variable(s), as well as pre-to-post resting baselines or contrasts | Pg. 29-30 |
| | 5c | Statistically compare the experimental condition/group to the control condition(s)/group(s) (not only each group to baseline measures) | Pg. 11-12 |
| Behaviour | 6a | Include measures of clinical or behavioural significance, defined *a priori*, and describe whether they were reached | Pg. 12 |
| | 6b | Run correlational analyses between regulation success and behavioural outcomes | |
| **Data storage** | | | |
| | 7a | Upload all materials, analysis scripts, code, and raw data used for analyses, as well as final values, to an open access data repository, when feasible | Pg. 17 |
